# Nutrient sensing pathways regulating adult reproductive diapause in *C. elegans*

**DOI:** 10.1101/2020.05.21.108340

**Authors:** Moriah Eustice, Jeff M. Reece, Daniel Konzman, Salil Ghosh, Jhullian Alston, Tyler Hansen, Andy Golden, Michelle R. Bond, John A. Hanover

## Abstract

Genetic and environmental manipulations, such as dietary restriction (DR), can improve both health span and lifespan in a wide range of organisms, including humans. Changes in nutrient intake trigger often overlapping metabolic pathways that can generate distinct or even opposite outputs depending on several factors, such as when DR occurs in the lifecycle of the organism or the nature of the changes in nutrients. Due to the complexity of metabolic pathways and the diversity in outputs, the underlying mechanisms regulating diet-associated pro-longevity are not yet well understood. Adult reproductive diapause (ARD) in the model organism *Caenorhabditis elegans* is a DR model that is associated with lengthened lifespan and reproductive potential (Angelo and Van Gilst 2009). As the metabolic pathways regulating ARD have not yet been explored in depth, we performed a candidate-based genetic screen analyzing select nutrient-sensing pathways to determine their contribution to the regulation of ARD. Focusing on the three phases of ARD (initiation, maintenance, and recovery), we find that ARD initiation is regulated by fatty acid metabolism, sirtuins, AMPK, and the O-linked N-acetyl glucosamine (O-GlcNAc) pathway. Although ARD maintenance was not significantly influenced by the nutrient sensors in our screen, we found that ARD recovery was modulated by energy sensing, stress response, insulin-like signaling, and the TOR pathway. We also discovered that fatty acid β-oxidation regulates ARD initiation through a pathway involving the O-GlcNAc cycling enzyme, OGT-1, acting with the nuclear hormone receptor NHR-49. Consistent with these findings, our analysis revealed a change in levels of neutral lipids associated with ARD entry defects. Our findings thus identify novel conserved genetic pathways required for ARD entry and recovery and identify new genetic interactions that provide insight into the role of OGT and OGA.

## INTRODUCTION

Aging is a leading risk factor for chronic human diseases such as diabetes, cancer, and neurodegeneration (Niccoli and Partridge 2012). Our understanding of aging has undergone a paradigm shift in the past 30 years as research indicates that aging may not be just an inevitable decline in health accompanied by increased disease incidence (Gems and Partridge 2013). Indeed, manipulation of genetic pathways and environmental conditions can modulate healthy aging (health span) and overall lifespan (Kenyon *et al.* 1993; Kenyon 2010). Thus, the aging process is now seen as a potential therapeutic target to improve health span in a rapidly growing elderly population (de Magalhães *et al.* 2012).

Dietary restriction (DR) is one of the most well-studied methods for increasing longevity in model organisms and has also been observed to improve overall markers of health in humans (Stein *et al.* 2012; Tinsley and La Bounty 2015). Specific changes in nutrient intake and the subsequent alteration to key metabolic processes is thought to contribute to the improvements in health and lifespan observed in a plethora of organisms (Fontana and Partridge 2015; López-Otín *et al.* 2016). However, identifying the underlying mechanisms and their role in longevity remains an intense and controversial area of research (Kennedy *et al.* 2007; Mair and Dillin 2008). While different forms of DR can trigger divergent pro-longevity pathways with distinct outcomes (Greer and Brunet 2009), they often share common features including involvement of overlapping nutrient-sensing factors and changes to metabolism, particularly fat and carbohydrate stores (Houtkooper *et al.* 2010; Lapierre and Hansen 2012; Finkel 2015; Solon-Biet *et al.* 2015).

DR-dependent pro-longevity pathways are highly conserved, suggesting research in model organisms, such as the nematode *Caenorhabditis elegans*, can inform dietary interventions and therapeutic targets relevant for improving human health span. For any organism, there is a delicate balance between nutrient acquisition, development, aging, and reproduction to ensure the health and propagation of the species. *C. elegans* is an exceptional model that has been used extensively to examine the intersection of longevity and these biological processes because it has a short life cycle, is highly fecund, and is genetically amendable with signaling pathways that are evolutionarily conserved.

Changes in nutrient status affect the life cycle of *C. elegans* at several key points during development. Extended periods with limited or no nutrients can result in a diapause state, after which the animal has a full or extended lifespan upon re-feeding (Hu 2007). In the more recently identified adult reproductive diapause (ARD) (Angelo and Van Gilst 2009), when the nematode undergoes DR in the mid-L4 larval stage, approximately 1/3 of worms remain in L4 (do not reach adulthood), 1/3 undergo bagging (matricide), and 1/3 enter the ARD state. ARD is characterized by worms that enter into adulthood with the retention of one to two “frozen” embryos, which are maintained throughout the starvation period, and the shrinkage of the germline to a small pool of germ cells. This pool of germ cells are capable of regenerating the entire germline after re-feeding and exit from ARD. A related study defined a process called the oogenic germline starvation response, in which germline shrinkage is linked to maintenance of the oocyte pool (Seidel and Kimble 2011). More recent findings also suggest that the transgenerational sterility observed in Piwi mutants defective in piRNAs may represent a dynamic form of ARD (Heestand *et al.* 2018). Thus, ARD represents a powerful system to analyze the interplay between DR, nutrient-sensing, germline dynamics, and lifespan extension.

A number of nutrient sensors have been found to play important roles under various DR conditions, including key factors in the target of rapamycin (TOR) pathway, AMP-activated protein kinase (AMPK) pathway, insulin signaling, hexosamine signaling, and fat metabolism pathway. Teasing apart the regulation of pro-longevity pathways has proven to be complicated, due in part to the complex web of interactions inherent in metabolic processes. For example, the transcription factor DAF-16/FoxO, which has been extensively researched for its role in pro-longevity, can be regulated by insulin-like signaling, TOR pathway, and sirtuin activity and, in turn, regulates transcription of components of the TOR pathway and the energy-sensing AMPK, among many others (Berdichevsky *et al.* 2006; Lant *et al.* 2010; Lapierre and Hansen 2012). In addition, numerous proteins required for pro-longevity pathways, such as various components of the insulin receptor signaling pathway, including FoxO (Housley *et al.* 2008; Yang *et al*. 2008; Klein *et al.* 2009), and AMPK (Bullen *et al.* 2014) are substrates of the enzyme O*-*GlcNAc transferase (OGT) and are post-translationally modified by O-GlcNAc, a downstream component of the hexosamine signaling pathway (Hanover *et al.* 2009). Further, other nutrient-sensing cellular components, including many known to play important roles in longevity, also interact physically and/or genetically with OGT and O-GlcNAcase (OGA), the enzyme responsible for O-GlcNAc removal (Ruan *et al.* 2013; Hardivillé and Hart 2014). While OGT and OGA are at the hub of many of the metabolic pathways important to pro-longevity they have not yet been analyzed extensively for their interactions and functions in longevity, in part due to their being essential in most organisms. However, OGT-1 and OGA-1 are not essential in *C. elegans* (Hanover *et al.* 2005; Forsythe *et al*. 2006), offering a unique opportunity to explore genetic interactions and analyze their role in longevity.

We sought to determine how nutrient status is sensed and transduced, what metabolic changes occur, and what genes are essential during the three distinct ARD phases. To explore the contribution of select nutrient sensors during ARD, we examined the ability of *C. elegans* strains to enter, maintain, and recover from ARD using mutants in key pathways that have well-characterized roles in longevity. Using a candidate gene approach, we examined key nodes in fat metabolism, insulin signaling, TOR, AMPK, sirtuin, and O-GlcNAc cycling (hexosamine) pathways. Additional epistasis studies revealed an important role the *nhr-49-*dependent fatty acid β-oxidation pathway for regulation of ARD entry in concert with O-GlcNAc cycling. These results have important implications for understanding the dynamic activity of nutrient sensors in longevity as well as furthering our understanding of the important functions and genetic interactions for OGT and OGA.

## MATERIALS AND METHODS

### Data Availability Statement

Supplemental figures, tables, methods and datasets have been uploaded via the GSA Figshare portal. These files include all of the supplemental figures (Supplmental Figures) mentioned in the text and table of raw data and statistical analysis (Supplemental Tables). Methods for quantification are also listed: https://gsajournals.figshare.com/s/d16c34f3167aac98f9e2

### Strains

*C. elegans* strains were maintained and cultured under standard laboratory conditions at 20°C with the food source *E. coli* OP50 as described (Brenner 1974). Specific alleles of *ogt-1* and *oga-1* used in this study have previously been characterized, including *ogt-1(ok1474), ogt-1(ok430)* (Hanover *et al.* 2005), *oga-1*(*tm3642*) (Bond *et al.* 2014), and *oga-1(ok1207)* (Forsythe *et al.* 2006). The alleles *ogt-1(jah01), oga-1(av82)*, and *oga-1(av83)* were generated during the course of this study using CRISPR/Cas9, as described below. The strain *daf-16(mu86)* was a gift from Dr. Michael Krause. The allele *nhr-49(nr2041)* was kindly gifted to us by Dr. Mark Van Gilst. The strains RB754 *aak-2(ok524)*, RB1988 *acs-2(ok2457)*, TJ1052 *age-1(hx546)*, BX153 *fat-7(wa36)*, RB1206 *rsks-1(ok1255)*, VC199 *sir-2.1(ok434)*, and QV225 *skn-1(zj15)* were obtained from the *Caenorhabditis* Genetics Center (CGC, University of Minnesota, Minneapolis, MN) strain bank, which is funded by the NIH Office of Research Infrastructure Programs (P40 OD010440). All of these strains were backcrossed to our wild-type N2 Bristol laboratory strain a minimum of four times prior to analysis to ensure homogenous genetic background.

Genotyping was conducted using nested PCR, as described by the CGC, or using sequencing (Eurofins Genomics, Louisville, KY) in the case of point mutants. Primers for *ogt-1(ok1474), ogt-1(ok430), oga-1(tm3642)*, and *oga-1(ok1207)* are described elsewhere (Bond *et al.* 2014, Forsythe *et al.* 2006) and are available upon request. Primers for *aak-2(ok524), acs-2(ok2457), rsks-1(ok1255)*, and *sir-2.1(ok434)* were derived from the sequences available on wormbase.org and CGC. Primers for *daf-16(mu86), nhr-49(nr2041)*, and the point mutants *age-1(hx546), fat-7(wa36)*, and *skn-1(zj15)* were generated using Primer3 (Rozen and Skaletsky 2000) and are available on request.

Double mutants were generated by crossing, with the presence of deletion alleles verified either using nested PCR or by sequencing, in the case of point mutants. Double mutants were backcrossed a minimum of four times to our wild-type N2 Bristol strain before use in analysis.

### Generation of CRISPR/Cas9 whole gene knockouts of *ogt-1* and *oga-1: ogt-1(jah01), oga-1(av82)*, and *oga-1(av83)*

The alleles *ogt-1(jah01), oga-1(av82)*, and *oga-1(av83)* were generated during the course of this study using CRISPR/Cas9 genome editing. These alleles are full gene deletions of the ORF of *ogt-1* and *oga-1*, respectively. We used the direct-delivery protocol developed by the Seydoux lab (Paix *et al.* 2017) and screened for edits using the co-conversion *dpy-10* method (Arribere *et al.* 2014). The crRNAs and repair template sequences are listed below. The repair template contains the deletion and perturbs the PAM sites to prevent Cas9 from re-cutting. From 15 injected P0s, we identified 3 “jackpots” and picked a total of 40 rollers from them. Using PCR genotyping, we identified and isolated three heterozygous lines from each. Homozygous animals were identified in the F2 generation and subjected to sequencing of the relevant *ogt-1* or *oga-1* region. For *ogt-1*, two lines contained the expected full deletion, which we subsequently named *ogt-1(jah01).* With *oga-1*, two of the three lines contained the expected deletion, while the third contained an additional ∼17bp deletion. The two expected *oga-1* deletions were named *oga-1(av82)*, which were the strains used in our analysis. We also kept the larger *oga-1* deletion and named it *oga-1(av83).*

### *ogt-1* sequences

Targeted gene crRNA1 (*ogt-1* deletion N-terminal)

5’ – AAUUUCAGAAUUAUAUCGAAGUUUUAGAGCUAUGCUGUUUUG – 3’

Targeted gene crRNA2 (*ogt-1* deletion C-terminal)

5’ – GCUUGUGAAUAGAUUUUCGAGUUUUAGAGCUAUGCUGUUUUG – 3’

Repair template:

5’ – CCAATAGTTCAATTTCCAAATTTCAGAATTATATCACACGGCTTGTGAATAGATTTTCGAAGGATTTTTA – 3’

### *oga-1* sequences

crRNA1 (*oga-1* deletion N*-*terminal):

5’ – GTTCACAGTCAATACTTAAT – 3’

crRNA2 (*oga-1* deletion C-terminal:

5’ – CAAAATCGTTGAATTTGACT – 3’

Repair template:

5’ – TTTTATTAAGAGTGAAAGTTGATGACAACATTTCCTGATTCAAATTCAACGATTTTGTTCCAACCGGGATAT – 3’

### Adult reproductive diapause

Adult reproductive diapause experiments were performed as described previously (Angelo and Van Gilst 2009), with minor adjustments. The overall experimental approach for initiation of ARD is illustrated in **Supplemental Figure 1**. Briefly, nematodes were cultured at 20°C on nematode growth medium (NGM) with *E. coli* OP50 as the food source at a density of 3,000 per 60 mm tissue culture dish. Worms were synchronized using hypochlorite bleaching of gravid worms with a dilute alkaline hypochlorite solution, as previously described (Porta-de-la-Riva *et al.* 2012). Eggs harvested from bleaching of gravid worms were then left with gentle rocking overnight at room temperature in M9 solution. Hatched L4s were then plated on NGM plus OP50 plates to complete synchronization. This synchronization process was repeated three times.

Following the third synchronization, 10,000 L1 worms (plate counting visually by sectors) were grown on a single 100 mm NGM + OP50 plate and visually inspected at intervals to ensure that over 90% of worms were in mid-L4 (approximately 48 hr, depending on growth of strain). Mid-L4 worms were then washed off the plate with M9 and collected in a 15 ml conical tube. The tubes were spun down at 20°C for 2 min at 1500 rpm. This washing was repeated 5 times with 5 ml M9 to ensure all bacteria was removed. Prior to the fifth and final spin, worms were incubated in 5 ml M9 with gentle rocking at room temperature for 30 min to remove bacteria from gut. Following the 30 min incubation, all media was aspirated and the nematodes were suspended in 1 ml of M9, counted, and plated on high agarose tissue culture plates without bacteria/food source (ARD plates). ARD plates were prepared and poured as follows: 1 L: 3 g NaCl and 15 g agarose autoclaved in 1 L water. After autoclave sterilization, filter sterilized solutions of the following were added: 1 ml of 1M CaCl_2_, 1 ml of 1M MgSO_4_, 25 ml potassium phosphate buffer (pH 6.0), and 1 ml cholesterol (8 mg/ml) (Angelo and Van Gilst 2009). Using this solution, 60 mm plates were filled to two-thirds full in the hood to ensure sterility. After plates were cooled and equalized to 20°C worms were added at a density of 3,000 worms per 60 mm plate. As a control, 3,000 of the synchronized and washed mid-L4s were plated separately from the worms that were put on the ARD plates and were instead plated on NGM plates with OP50 to allow analysis of lifespan and brood size in parallel to starved worms used to assess the ARD phenotype.

Worms were visually inspected every 24 hours (up to 72 hours, depending on strain) after plating and characterized as either in ARD, in L4, or bagged. At day 5, day 10, and day 30 after mid-L4 worms were plated on ARD plates, worms were subjected to Oil Red O (ORO) staining, carminic acid staining, lifespan analysis, and brood size counts as described below. Each experiment was repeated a minimum of three times from start to finish.

### Lifespan, germline, and brood analysis

At least 10 worms from each ARD plate were collected individually at day 5, day 10, and day 30 post-plating on ARD plates. The worms were visually inspected to ensure ARD status and placed singly onto individual NGM plus OP50 plates and incubated at 20°C to monitor for restoration of the germline, analysis of overall number of progeny following ARD (brood size), and post-ARD lifespan. Restoration of the germline was followed visually at day 5, day 10, and day 30 using standard microscopy techniques on a Zeiss LSM 700 confocal microscope (Carl Zeiss Microscopy, LLC, Thornwood, NY) with a Plan-Apochromat 20x/0.8 objective lens. Brood size was monitored by carefully transferring the individually recovered worms every 24 hr to fresh NGM plus OP50 plates until the end of their reproductive phase, marked by the absence of eggs. Progeny of the post-ARD worms were counted after each transfer and added to total brood size reported for each animal. For all experiments, any ARD worm that failed to thrive in the first 48 hr or disappeared was censored.

The post-ARD (recovered) worms were then inspected daily after the reproductive period to determine lifespan. When worms no longer responded to gentle poking with a platinum pick, they were scored as dead. Lifespan counts for post-ARD animals includes the days in ARD, such that the lifespan post-re-feeding was added to the number of days the animals were starved. Lifespan was graphed, and statistics were performed as previously described using GraphPad and one-way ANOVA (Bond *et al*., 2014). A minimum of 40 ARD animals from four independent experiments were used for analysis of each strain.

The controls also underwent the same analysis but without ever being placed on the ARD plates and, instead, were grown on standard NGM plus OP50 plates at 20°C.

### Oil Red O staining

The protocol for Oil Red O (ORO) staining was adapted from previously described methods (O’Rourke *et al.* 2009) with minor adjustments. Prior to starting experiment, a stock solution of 0.5 g/100 ml ORO in isopropanol was allowed to equilibrate for at least two days at room temperature with gentle rocking and light protection. On the day of the experiment, a fresh dilution of ORO stock solution was diluted to 60% with water and rocked for at least 1 hr. Dilute ORO was then filter sterilized with a 0.22 µm filter.

Nematodes were washed off NGM plates with OP50 (control) or high agarose ARD plates with 1 ml of 1x PBS + 0.01% Triton X-100 (PBST). Worms were washed 3 times with 1 ml of PBST and allowed to settle by gravity after each washing. Cuticles were permeabilized by resuspending worms in 120 μl of PBST and 120 μl of 2x MRWB buffer (160 mM KCl, 40 mM NaCl, 14 mM Na_2_EGTA, 1 mM spermidine-HCl, 0.4 mM spermine, 30 mM Na-PIPES pH 7.4, and 0.2% β-mercaptoethanol) with 2% paraformaldehyde. Worms were gently rocked for 1 hr at room temperature. Animals were allowed to settle by gravity, buffer was aspirated, and animals were washed with 1x PBST twice. Worms were then resuspended in 60% isopropanol and incubated 15 min with gentle rocking at room temperature to dehydrate. Worms were allowed to settle by gravity and remaining isopropanol was removed. 1 ml of 60% ORO was added and the worms were incubated overnight with gentle rocking. After the overnight incubation, ORO was removed and worms were resuspended in 100 μl of 1x PBST. 1 μl 1x DAPI was then added to the 20 μl PBST, as an internal control for permeabilization, and flicked to mix. Worms were then mounted on slides on an agarose pad and visualized with Zeiss LSM 700 confocal microscope using either a Plan-Apochromat 10x/0.45 or Plan-Apochromat 20x/0.8 objective lens, as described below.

### Carminic acid staining

Carmine dyes such as carminic acid are used to complex with glycogen and short glucose polymers to produce a fluorescent signal responsive to the levels of glucose polymer. Thus, it is a useful tool to analyze glycogen storage (Hanover *et al.* 2005). Carminic acid was introduced to worms through feeding as previously described (Hanover *et al*. 2005). Briefly, OP50 was inoculated into LB broth with 1 mg/ml carminic acid and grown overnight at room temperature in a light protected vial. OP50 with carminic acid was then used to seed standard NGM plates. Animals were plated on carminic acid containing plates following initial mid-L4 harvest (controls) or post-ARD at day 5, day 10, and day 30 and were visualized with Zeiss LSM 700 confocal microscope using a Plan-Apochromat 20x/0.8 objective lens, as detailed below.

### Microscopy & Image Analysis

Microscope images were acquired on a Zeiss LSM 700 confocal microscope using either Plan-Apochromat 10x/0.45 or a Plan-Apochromat 20x/0.8 objective lens. In all cases: photomultiplier tube (PMT) offset was adjusted so that zero photons corresponded to a pixel value of zero; and PMT gain was adjusted to fully utilize the available 8-bit dynamic range while avoiding saturating the images. The same settings were used across all images being compared.

When acquiring the (non-fluorescence) ORO images, sequential scans were performed with the 488nm, 555nm, and 639nm lasers while using the transmitted light detector to produce an RGB image like what one would acquire using a color camera under widefield illumination from white light. Differential Interference Contrast (DIC) optics were also in place while imaging to provide contrast sufficient to distinguish sample anatomy at the same time as recording ORO. Either the 10x/0.45 or the 20x/.8 objective lens was used for ORO imaging— there was no difference in quantitation because the numerical aperture of the condenser was limited to <0.45, and kept constant across experiments.

When acquiring fluorescence images of carminic acid, a 555nm laser was used for excitation. The emission was passed through a laser blocking filter and a wide open pinhole to produce a non-confocal image. The emission was further filtered through a SP640 (640 nm short pass) emission filter, to produce a widefield fluorescence image appropriate for quantitating total signal from the sample volume. Simultaneously with the fluorescence channel, a DIC image was also acquired. Only images acquired with the 20x/0.8 lens were used for quantitation.

For quantitative image analysis, Fiji software (Schindelin *et al*. 2012) was used, employing in-house macros customized to streamline the processing for the user. For both ORO and carminic acid, regions of interest (ROIs) were manually traced very roughly around the edges of each worm, further refined by automatic segmentation, then the average intensity per pixel was calculated. This mean pixel value was divided by the diameter of the worm, measured manually by the user, to provide a relative value for quantitating mean carminic acid fluorescence per unit volume. For the ORO (RGB) images, a more in-depth analysis was performed, converting the intensities to relative absorbance (pseudo-fluorescence) at each pixel location, then combining with automated physical measurements of the worm from the ROI, to calculate an average ORO density per unit volume. Quantitative comparisons could then be made (see **Supplementary Methods**). For presentation, all images had their intensities re-scaled to maximize the dynamic range and visual appeal of the images. Images of ORO and fluorescence that were being compared were rescaled by the same amount in those channels.

## RESULTS

### Diverse nutrient sensors are required for ARD entry

To date, only the *nhr-49(nr2041)* mutant allele of NHR-49, a protein involved in fatty acid metabolism, has been characterized as having a significant role in ARD entry (Angelo and Van Gilst 2009). We therefore sought to identify additional regulators of ARD dynamics, focusing on the three distinct phases of ARD; initiation (number of animals entering ARD vs bagging or L4 fate), maintenance (germline shrinkage and embryo arrest), and recovery (post-ARD lifespan, germline regrowth, and brood size) (Angelo and Van Gilst 2009) (**Figure 1A**). Using a candidate gene approach, we focused on key nutrient-responsive pathways. The pathways profiled included AMP/energy signaling (AAK-2/AMPK), insulin-like signaling (AGE*-*1/PI(3)K, DAF-16/FoxO), TOR (RSKS-1/S6K), stress signaling (SKN-1/Nrf), carbohydrate metabolism/hexosamine signaling (OGT-1/OGT and OGA-1/OGA), and sirtuins (SIR-2.1/Sirt1). The alleles analyzed were chosen based on their viability under the experimental conditions and that they have been previously characterized for their roles in longevity. Note that the raw data for all experiments described herein is available in the **Supplementary Tables.**

**Figure 1.**
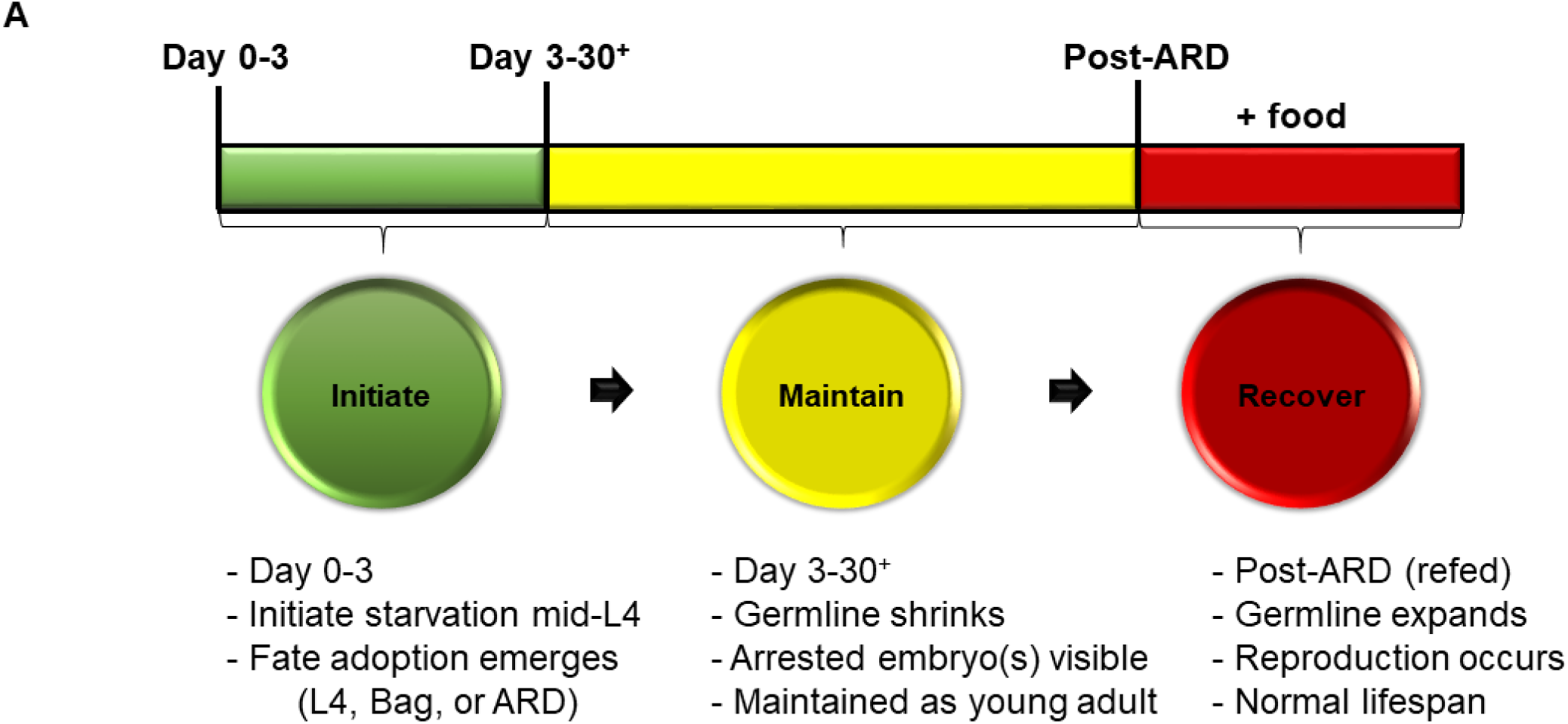
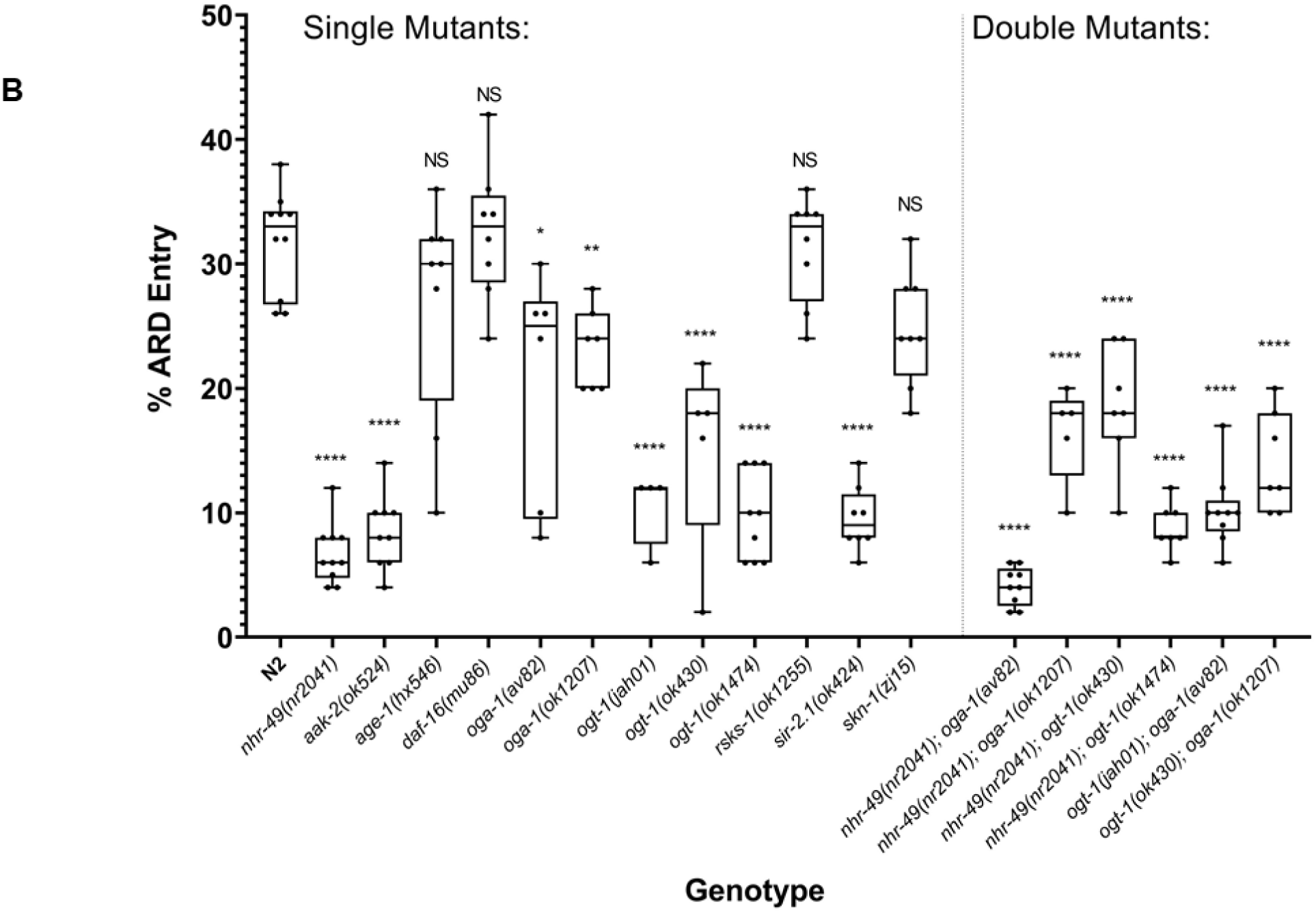
ARD stages and entry defects in nutrient sensing mutants. (A) ARD stages including initiation, remodeling/maintenance, and recovery. Initiation occurs when worms are removed from a food source during mid-L4. With wild type worms, approximately 1/3 enter ARD, 1/3 have hatched offspring inside (bagging), and 1/3 maintain an L4 state during the first 72 hours of starvation. Once ARD is initiated, worms enter the maintenance phase wherein the germline shrinks and the characteristic arrested embryo(s) are observed. Recovery occurs after starved worms are re-fed. During this stage the germline expands and a normal lifespan commences (adapted from Angelo and Van Gilst 2009). (B) For single mutants ARD initiation was significantly defective in the following strains, as compared to N2: *nhr-49(nr2041*), *aak-2(ok524), oga-1(av82), oga-(ok1207), ogt-1(jah01), ogt-1(ok430), ogt-1(ok1474)*, and *sir-2.1(ok424)*. For epistatic analysis, the double mutants *nhr-49(nr2041);oga-1(av82)* and *nhr-49(nr2041);oga-1(ok1207)* had phenotypes most similar to *nhr-49(nr2041).* The double mutants *nhr-49(nr2041);ogt-1(ok430)* and *nhr-49(2041);ogt-1(ok1474)* were most similar to the single *ogt-1* mutants. The entry defects for double mutants of *ogt-1;oga-1* alleles were similar to *ogt-1* single mutants. A minimum of 300 worms from 3 different experiments were counted and statistics were performed in Graph Pad Prism using an Ordinary one-way ANOVA. Statistics are summarized in **Supplemental Table 1**. *P-*value **** = <0.0001, ** = 0.001, * = 0.0265.

ARD initiation is clearly discernible by 48 to 72 hours after mid-L4 worms are placed on ARD plates and can be distinguished from the bagging or L4 fates by the morphology of the worms, particularly the presence of one to two “frozen” embryos in adult worms (Angelo and Van Gilst 2009) (**Supplemental Figure 1**). We found that *age-1(hx546), daf-16(mu86), rsks-1(ok1255)*, and *skn-1(zj15)* did not significantly alter ARD initiation, suggesting that insulin-like signaling, the TOR pathway, and stress response are not major contributors to entry (**Figure 1B**). In contrast, *aak-2(ok524), oga-1(ok1207), ogt-1(ok430), ogt-1(ok1474)* (including additional alleles of *oga-1* and *ogt-1*, discussed below), *sir-2.1(ok434)*, and our positive control *nhr-49(nr2041)* all contribute to the initial sensing and response to starvation during ARD entry, as these mutants displayed a significant reduction in the ability of nematodes to successfully initiate ARD (**Figure 1B, Supplemental Figure 2**).

Associated with ARD initiation, we noted significant differences between the mutant strains in the distribution of animals adopting either the L4 or bagging fate. Both L4 and bagging had a higher degree of variability than ARD entry, in general, suggesting that the ARD entry may be more tightly regulated than the other two fates (**Supplemental Figure 2; Supplemental Table 1**). Interestingly, some of the mutant alleles showing reduced ARD entry (*nhr-49(nr2041), ogt-1(ok1474)*, and *oga-1(ok1207)*) did so while producing sizable numbers of animals adopting both the L4 fate and bagging. The other strong entry defective alleles, *aak-2(ok524)* and *sir-2.1(ok434)*, produced predominantly L4 arrested larvae with few bagging or in ARD (**Supplemental Figure 2; Supplemental Table 1**), suggesting there may be alternative mechanisms at play in the regulation of entry between mutants and, therefore, distinct pathways.

We were intrigued to find that *oga-1* and *ogt-1* mutants exhibited severe ARD entry defects similar to *nhr-49(nr2041)* (**Figure 1B; Supplemental Figure 2; Supplemental Table 1**). We verified this novel finding by generating and analyzing targeted full gene deletions, *oga-1(av82)* and *ogt-1(jah01)*. These new CRISPR/Cas9 alleles were confirmed to have similar phenotypes as the previously characterized *ogt-1* and *oga-1* alleles, including lifespan and brood size under normal laboratory conditions (**Figure 3A; Supplemental Table 2**). As with the other two *ogt-1* alleles, we observed a highly significant ARD entry defect with *ogt-1(jah01)* (**Figure 1B**). The *oga-1* alleles tested all also showed significant reduction in ARD entry, but were more variable and showed a milder phenotype than the *ogt-1* alleles (**Figure 1B, Supplemental Figure 2**), suggesting the removal of O-GlcNAc does not have as strong of an influence as addition of O-GlcNAc.

**Figure 2.**
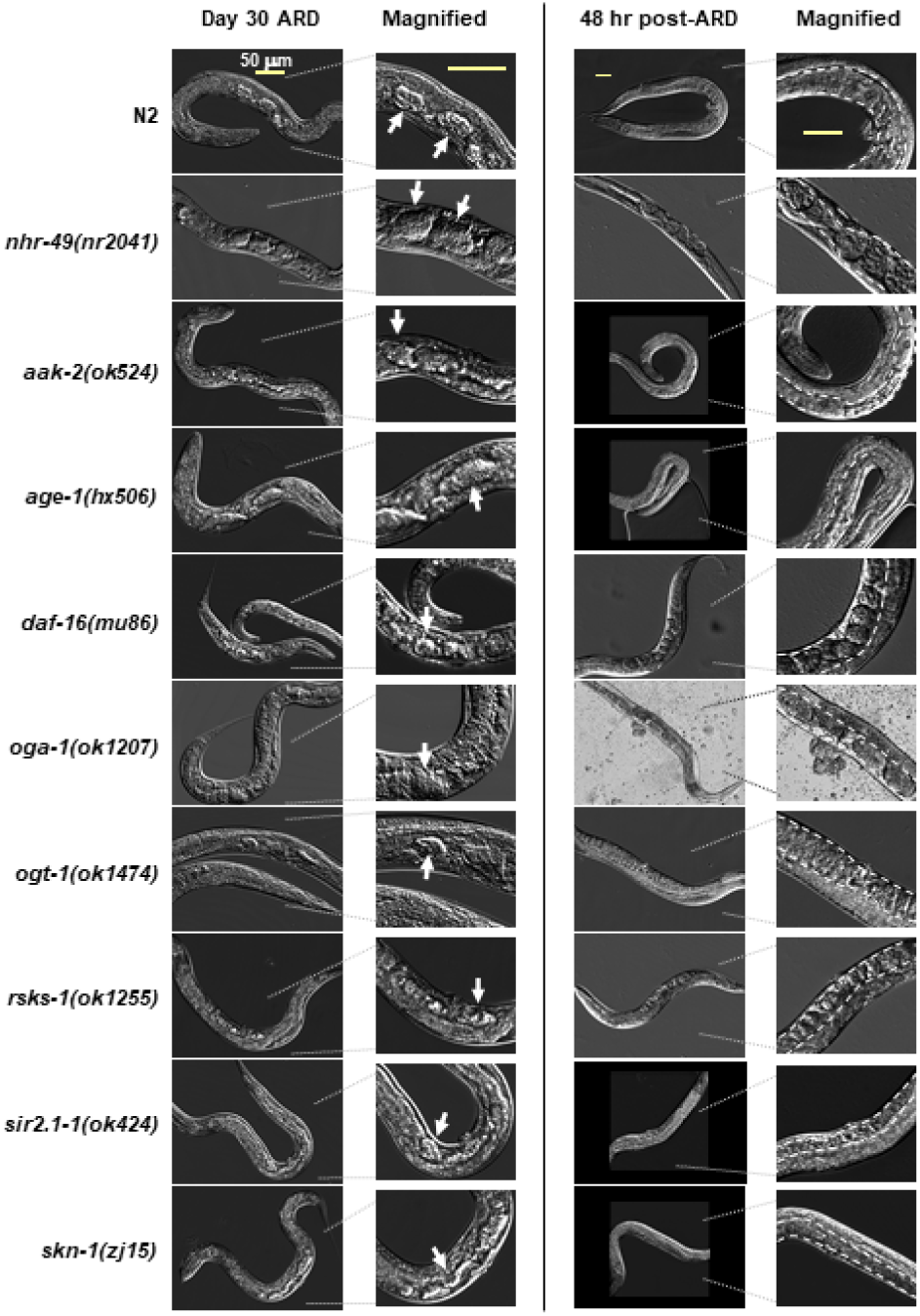
Germline shrinkage during ARD maintenance and regrowth during recovery. Representative images (with magnified panels for detail) showing the germline of worms starved (left panel) at day 30 of ARD, with arrested embryos noted with white arrows. The right panel shows worms that have been re-feed for 48 hours after 30 days of ARD, with white dashed lines to indicate the germline, where healthy embryos and/or oocytes can be seen. All strains showed robust shrinkage of the germline during starvation and regrowth after feeding. The 50 μm scale bar shown in the top row applies to all images in each column.

**Figure 3.**
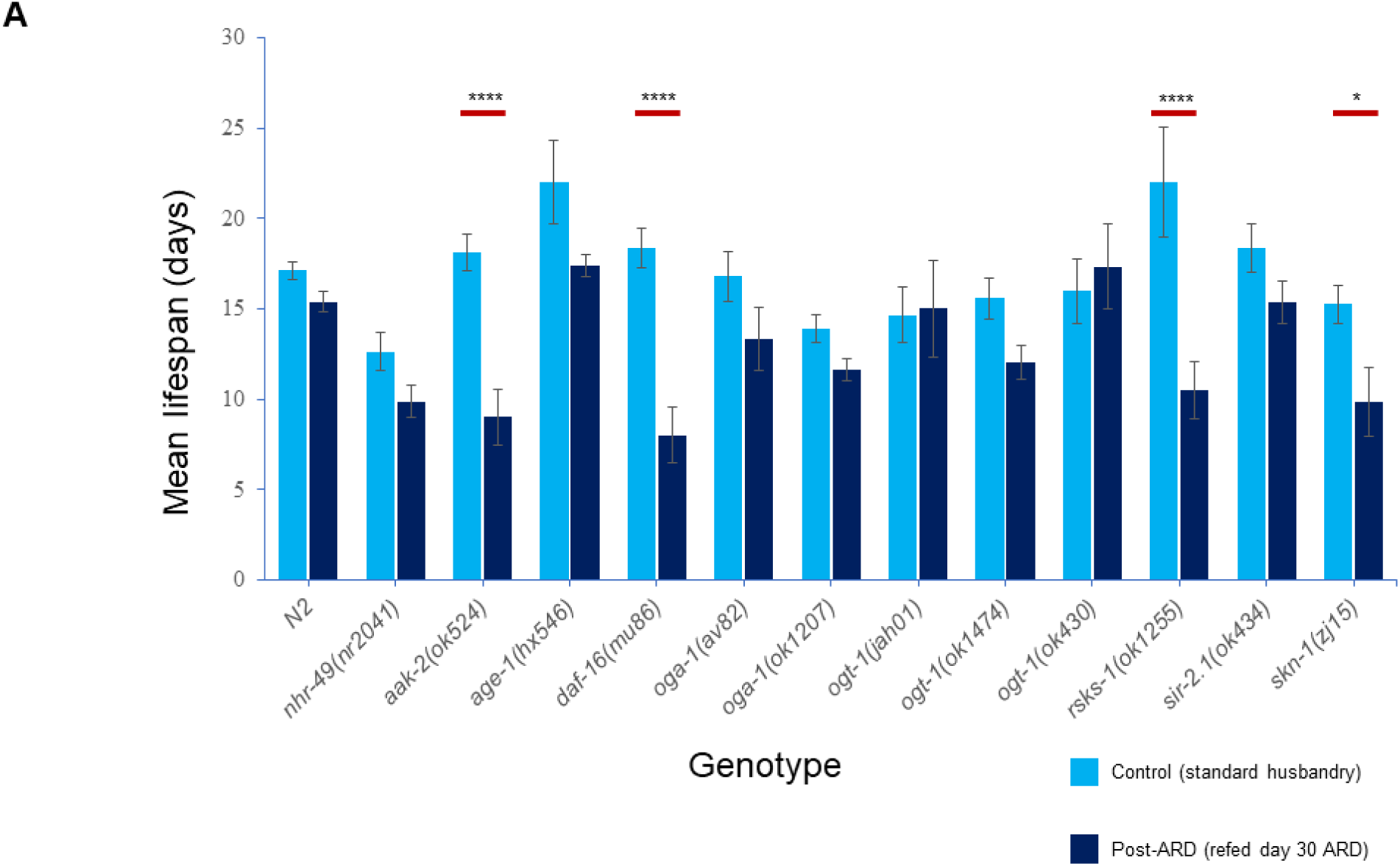
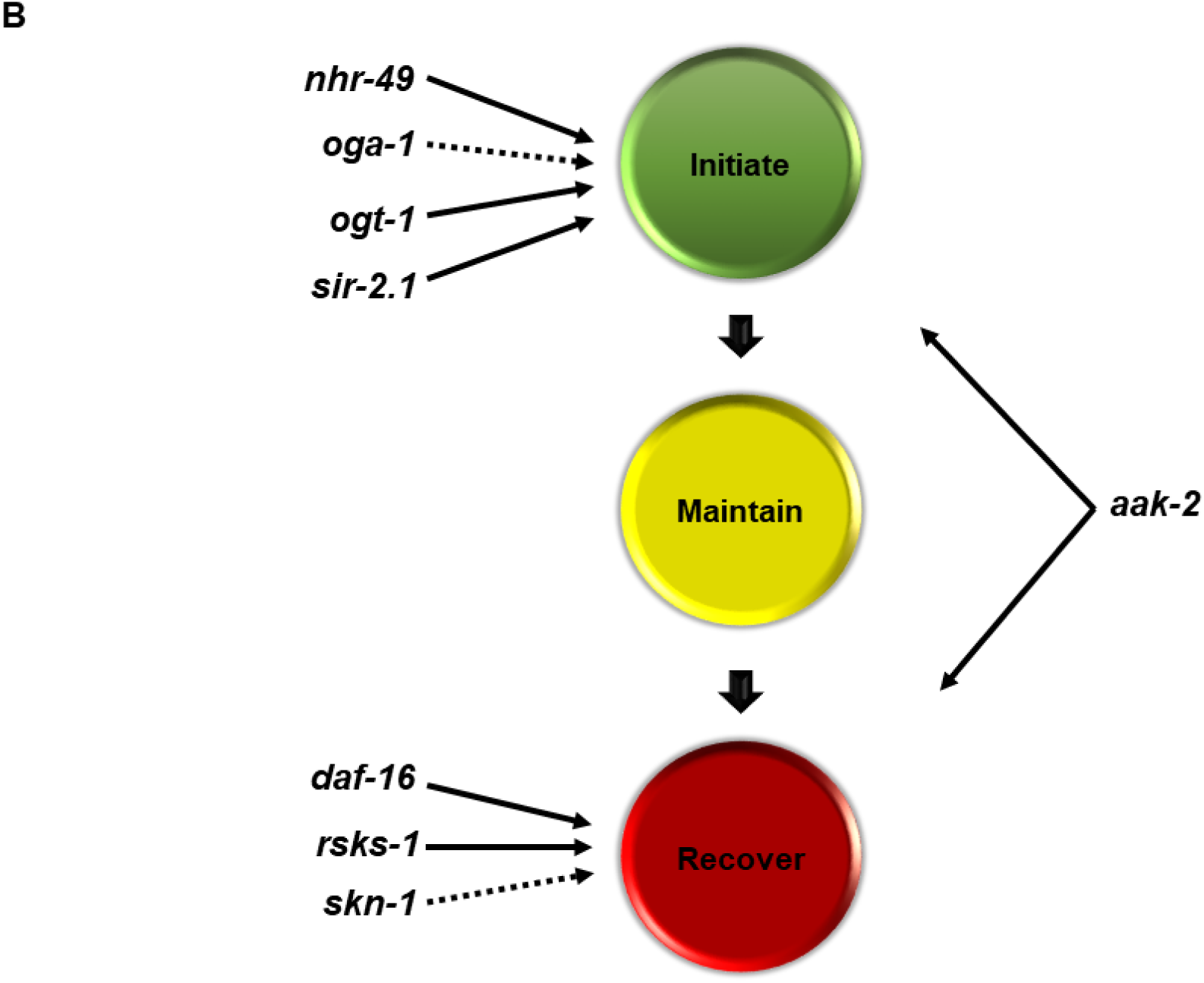
Lifespan of mutant strains in recovery after 30 days of ARD and summary model. (A) Lifespan of wild-type and mutant strains in control husbandry conditions compared to those recovered after 30 days of ARD. Lifespan after recovery from ARD was significantly defective in *aak-2(ok524), daf-16(mu86)*, and *rsks-1(ok1255).* The mutant *skn-1(zj15)* also had a significantly reduced lifespan post-ARD but not as strongly as the others. A minimum of 8 worms from independent experiments were analyzed for lifespan and statistics were performed in Graph Pad Prism using one-way ANOVA. *P-*value **** = <0.0001, * = 0.0448. (B) Summary of mutants altering ARD stages. While none of the mutants analyzed herein influenced maintenance to a notable degree, *nhr-49, ogt-1*, and *sir-2.1* all strongly influenced ARD entry, with *oga-1* also playing a role but to a lesser degree than the other three. The mutations in *daf-16* and *rsks-1* had a strong impact on recovery whereas *skn-1* had an influence but to a lesser degree. The *aak-2* mutants were the only strain that influenced both ARD initiation and maintanence.

### Genes encoding *O-*GlcNAc cycling enzymes work in concert with *NHR-49* to modulate ARD initiation

Based on the similarity of phenotypes for the single mutants of *nhr-49, oga-1*, and *ogt-1*, as well as their known overlapping roles in metabolism, we next chose to explore the genetic interaction between these mutants by analyzing the ARD initiation phenotypes in double mutants. Our epistasis analysis suggested that these three genes act in the same pathway. Based on action of the gene products, we place *ogt-1* upstream of both *nhr-49* and *oga-1*, and *oga-1* acting upstream of *nhr-49.* Double mutants with an *ogt-1* allele and either an *nhr-49* or *oga-1* allele had entry defects roughly equivelant to the *ogt-1* allele on its own. Compared to the strong defects in entry observed for *nhr-49* and *ogt-1* mutants, *oga-1* mutants had a more intermediate entry defect. Analysis of double mutants showed *oga-1* alleles behaved differently when with *nhr-49*, as *nhr-49(nr2041); oga-1(av82)* had a severe entry defect like *nhr-49*, while *nhr-49(nr2041); oga-1(ok1207)* had a more modest phenotype similar to other *oga-1* alleles (**Figure 1B**). Taken together, our data suggests that *nhr-49, oga-1*, and *ogt-1* function in the same pathway to regulate ARD entry.

### Distinct nutrient sensors are required for ARD recovery

Successful maintenance of ARD can be determined by examining germline shrinkage and embryo preservation that occurs after entry into ARD. Characteristics of the recovery stage of ARD is marked by the dynamic germline regrowth, production of progeny (brood size), and post-ARD lifespan (Angelo and Van Gilst 2009). Thus, to further define which nutrient sensors regulate each of these stages of ARD, we next looked at the dynamics of the germline, brood size, and lifespan.

Despite significant changes in ARD entry, we observed that all mutant strains exhibited robust germline shrinkage once ARD was established, analogous to wild-type worms and the frozen embryos were retained throughout the starvation period (**Figure 2, left panel**). Further, post-ARD, where worms were re-fed for two days following 30 days of starvation, a robust regrowth of the germline was observed in all strains (**Figure 2, right panel**). Thus, in this set of nutrient sensors, dynamic germline shrinkage and regrowth is a general feature of ARD once it is established, as is the maintenance of the “frozen” embryos. Notably, only the apoptosis factor CED-3 has been found to disrupt maintenance phenotypes to date (Angelo and Van Gilst 2009), suggesting that the maintenance of ARD may be independent of nutrient sensing and supporting the hypothesis that the distinct phases of ARD may be genetically separable (Angelo and Van Gilst 2009; see also **Discussion**).

As noted previously, post-ARD animals retain their ability to reproduce, albeit to a significantly diminished degree. However, this was found to be largely dependent on the amount of sperm retained in the hermaphrodites, with selfing producing significantly less offspring than mating post-ARD (Angelo and Van Gilst 2009). In our analysis of brood size following ARD, we found that there was a high level of variability in the number of progeny, even between replicates of the same strains within the same experiment (**Supplemental Table 2**). Surprisingly, reduced brood size did not correlate with reduced sperm counts (**Supplemental Figure 3**) nor with the germline dynamics we observed (**Figure 2**). This suggests that the relationship between ARD recovery as measured by brood size is complex and may not correlate to sensing and responding to nutrient changes but may involve other processes (see **Discussion**). We therefore chose to focus on post-ARD lifespan as a measure of ARD recovery.

Post-ARD lifespan is a means to quantify the survivability of ARD, or the ability to recover and thrive following re-feeding after time spent in ARD (Angelo and Van Gilst 2009). Thus, we analyzed the lifespan of worms recovered after 30 days of ARD using standard lifespan assays. We found that *aak-2(ok524), daf-16(mu86), rsks-1(ok1255)*, and, to a lesser extent, *skn-1(zj15)* mutants had significantly reduced post-ARD lifespans (**Figure 3A**). Based on the phenotypes of the different mutants during the various stages of ARD, we suggest that recovery may largely be regulated by the TOR pathway, as opposed to insulin-like signaling or stress response (see **Discussion**).

Overall, we found that whereas *nhr-49, oga-1, ogt-1*, and *sir-2.1* play a role in ARD initiation, *daf-16, rsks-1*, and *skn-1* function primarily in recovery from ARD, with *aak-2* playing a role in the regulation of both states. Conversely, none of the alleles that we explored significantly impacted maintenance of the ARD (summarized in **Figure 3B**).

### Mitochondrial β-oxidation but not fatty acid desaturation is required for ARD entry

Due to the strong phenotypes we observed for ARD entry, the novel role for O-GlcNAc signaling, and the similarity in phenotypes for *nhr-49* and *ogt-1* mutants, we next explored this entry pathway in more detail. We chose to focus of *ogt-1* and not *oga-1* for the rest of the experiments due to the variability of *oga-1* alleles and the weaker entry phenotype.

NHR-49 transcriptionally regulates both fatty acid desaturation, via FAT-7, and mitochondrial β-oxidation, via ACS-2 (Van Gilst *et al.* 2005a) (**Figure 4A**). To determine which of these NHR-49-dependent pathways is relevant for ARD entry we analyzed *acs-2(ok2457)* (mitochondrial beta-oxidation) and *fat-7(wa36)* (fatty acid desaturation) mutants for ARD phenotypes. Whereas the *acs-2* mutant had a significant defect in ARD entry, mutant *fat-7* entry was comparable to wild-type. Further, the double mutant *acs-2(ok2457);ogt-1(ok1474)* was most similar to the *ogt-1(ok1474)* single mutant phenotype for percentage of each fate adopted (**Figure 4B; Supplemental Figure 2**). Thus, our findings demonstrate that *acs-2* plays a role in the initiation of ARD entry downstream of *ogt-1*. As we had observed with the *nhr-49, oga-1*, and *ogt-1* single mutants, we did not observe a significant difference in lifespan of the *fat-7* or *acs-2* mutants post-ARD, suggesting that *acs-2* function in ARD entry is also separable from recovery (**Figure 4C**).

**Figure 4.**
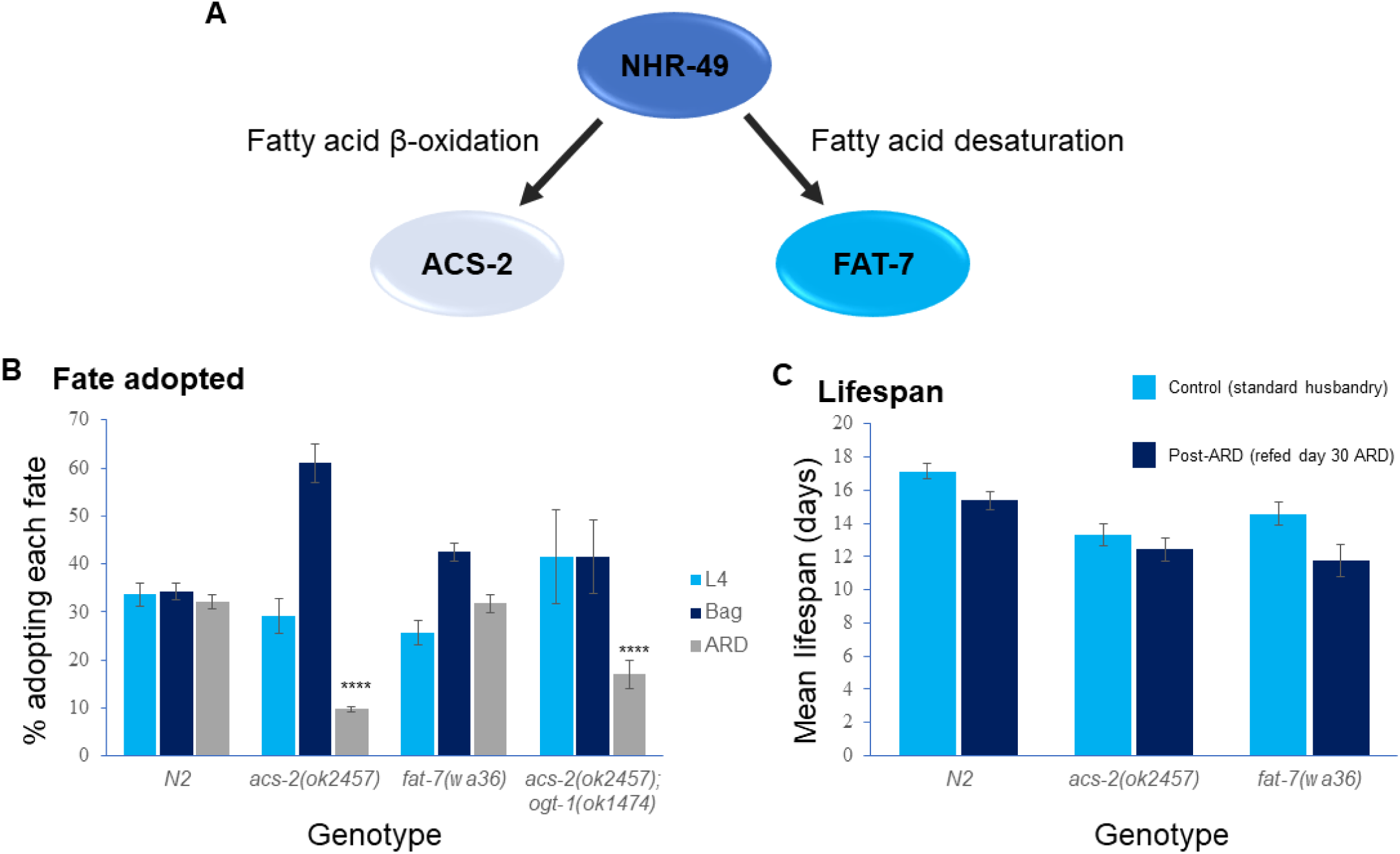
Fatty acid beta-oxidation, but not desaturation, plays a role in ARD entry. (A) NHR-49 regulates both fatty acid beta-oxidation (ACS-2) and desaturation (FAT-7) via genetically separable pathways that are known to generate distinct phenotypic outputs. (B) ARD entry was significantly impaired with *acs-2(ok2457)* but not with the *fat-7(wa86)* mutant. The double mutant *acs-1(ok2457); ogt-1(ok1474)* was also significantly impaired for ARD entry. *P-*value **** = <0.0001 in comparison with N2 as determined using one-way ANOVA. (C) Lifespan under standard conditions and with refeeding at day 30 of ARD was not significantly altered in these mutants as determined by one-way ANOVA.

### Neutral lipid changes associated with ARD entry

Changes in lipid metabolism play a prominent role in multiple pro-longevity pathways (Schroeder and Brunet 2015). NHR-49 and the O-GlcNAc cycling enzymes have well-established roles in the regulation of fatty acid metabolism (Van Gilst *et al.* 2005a; Van Gilst *et al.* 2005b; Bond and Hanover 2014); however, lipid storage in ARD has not yet been directly analyzed.

To determine if the genetic interaction we observed between *nhr-49* and *ogt-1* during ARD initiation was relevant to lipid metabolism, we used the lysochrome dye Oil Red O (ORO), which is widely used to stain triglycerides (TAGs) (O’Rourke *et al.* 2009). We examined the various strains for ORO staining in control (fed conditions) and after 30 days in ARD (prolonged exposure). Fed *ogt-1(ok1474)* worms showed significantly lower ORO staining compared to N2s (**Figure 5A**, quantified **Figure 5B**), though the distribution of signal was not noticeable different. ORO staining is present throughout the body of worms in fed conditions, but is greatly diminished and becomes largely restricted to arrested oocytes after 30 days in ARD (**Figure 5A**, quantified **Figure 5B**). This restriction of ORO staining to the frozen embryos was significantly stronger in the single mutant worms *nhr-49(nr2041)* and *ogt-1(ok1474)* worms, which also showed the lowest amounts of ORO staining overall (**Figure 5A**). Between the fed and ARD conditions, *ogt-1(ok1474)* animals did not show a significant decrease in quantified ORO staining, likely due to the already low lipid stores in this strain. Additionally, the quantification was normalized by volume, which may underestimate the effect of ARD on total triglycerides per animal, as worms in ARD are significantly smaller than fed worms.

**Figure 5.**
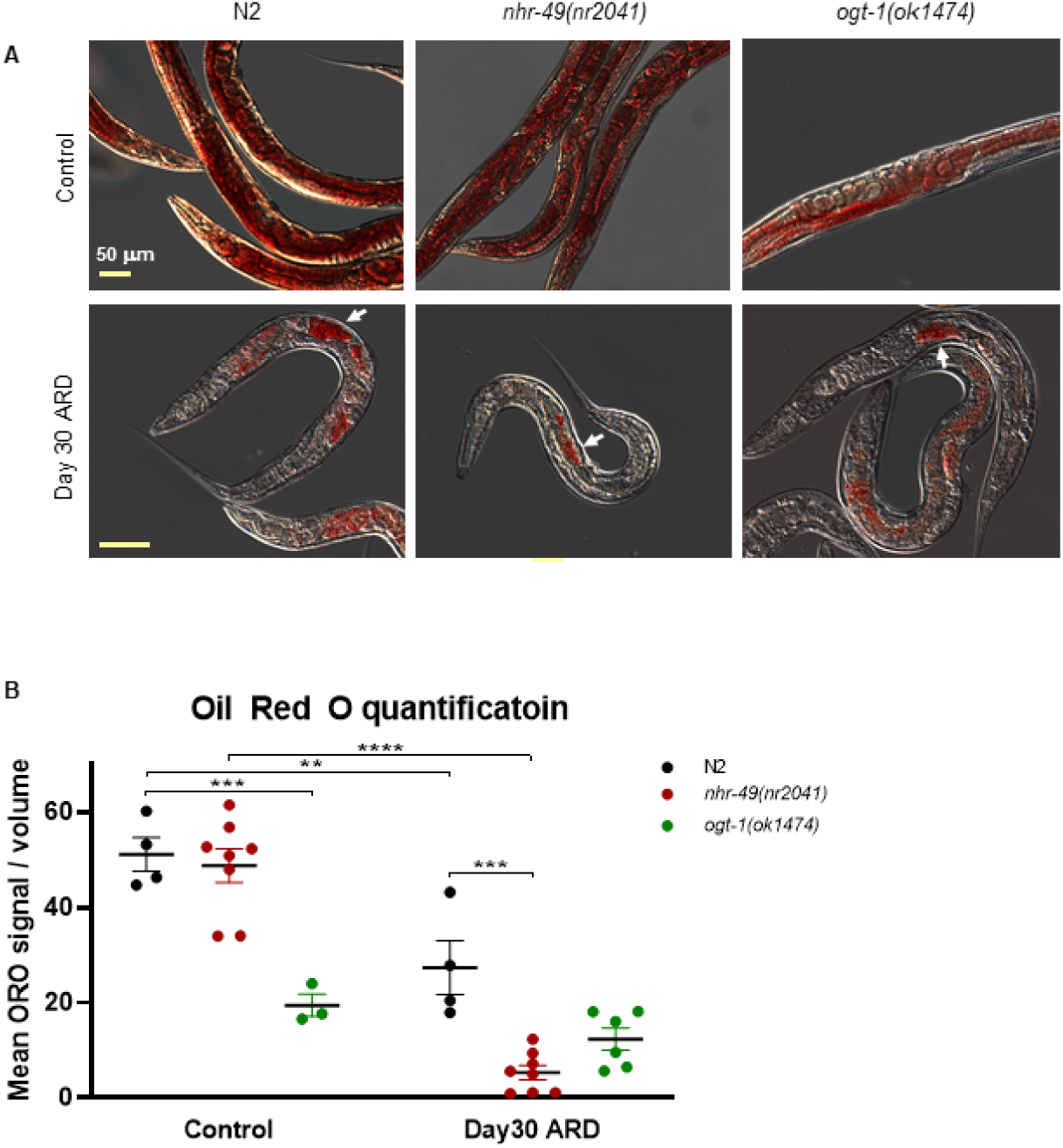
Oil Red O staining reveals redistribution of lipid stores in ARD. (A) Representative images showing the distribution of Oil Red O in control (top) and starved animals after day 30 of ARD (bottom) in the *ogt-1-*dependent pathway. Retention of TAGs was observed to be strongest in the arrested embryos in all mutant strains (white arrows). Scale bar indicates 50 μm and is consistent across each row. (B) Fiji Image analysis-based quantification of ORO staining (see **Supplemental Methods**) of wildtype and the mutant strains of the *ogt-1-*dependent pathway in normal husbandry (control) and in day 30 of ARD. Because ORO quantification was normalized by volume and ARD worms are significantly smaller than well-fed adults, the values shown likely underestimate reductions in lipid stores. Using two-way ANOVA analysis, both variables (genotype and control or ARD condition) were found to have significant effects, and these variables interact significantly. For each strain, mean ORO density decreased after 30 days of ARD: most dramatically for *nhr-49(nr2041)* and not significantly for *ogt-1(ok1474).* In normal husbandry conditions, *ogt-1(1474)* showed significantly less ORO signal than did N2 worms. *P-*value **** = <0.0001, *** = <0.001, ** = <0.01.

Notably, we observed the strong depletion of lipid stores as early as 24 hours of starvation (**Supplemental Figure 4A**), suggesting it is an early response. As a control, we also monitored cuticle penetrance of a small molecule dye using DAPI which revealed comparable penetrance between strains (**Supplemental Figure 4B**). We further observed that in other strains analyzed, those without major effects on ARD entry (*age-1(hx546), fat-7(wa36)*, and *skn-1(zj15)*) showed higher levels of ORO staining throughout the animal in ARD than wild-type worms in ARD (**Supplemental Figure 5, Figure 5**). Strains with significant ARD entry defects (*acs-2(ok2457), sir-2.1(ok424)*, and *oga-1(tm3642)*) showed reduced ORO staining that is more highly localized to the arrected embryo while in ARD (**Supplemental Figure 5**). This correlation suggests lipid stores play an important role in the initiation of this form of diapause.

Based on the interplay between carbohydrate and fatty acid metabolism, as well as the role of O-GlcNAc in carbohydrate metabolism, we looked next at glycogen storage using carminic acid staining. Under control husbandry conditions, our analysis suggested a change in the pattern of glycogen storage in *nhr-49(nr2041)* and *ogt-1(ok1474)* worms, though quantification of the fluorescent signal showed no significant difference between strains (**Supplemental Figure 6**). After 30 days of ARD, these stains all showed higher carminic acid signal normalized by volume. This may be patially attributable to the reduction in size associated with ARD, and suggests glycogen stores are relatively unaffected or increased by ARD. Interestingly, *nhr-49(nr2041)* worms in ARD showed dramaticly higher carminic acid staining (a 3-fold increase compared to N2) than did non-ARD worms, to an extent we did not observe with the other strains assessed (**Supplemental Figure 6**). With our findings of lipid reduction in ARD, this suggests glycogen stores may be maintained or increased at the expense of lipid stores while in ARD. These results suggest ARD has interesting effects on glycogen storage, but do not correlate with the entry defects we see in *nhr-49* and *ogt-1* mutant strains.

## DISCUSSION

### Diverse roles for nutrient sensors at different phases of ARD

Building on the previously discovered role of *nhr-49* in ARD entry (Angelo and Van Gilst 2009), we demonstrate that *aak-2, oga-1, ogt-1*, and *sir-2.1* also regulate ARD entry, thus implicating fat metabolism, the hexosamine pathway, and energy status in the initial sensing and dissemination of the ARD entry signal. Further, based on our genetic epistasis analysis and phenotypic similarities, we suggest that *nhr-49, oga-1*, and *ogt-1* act in the same pathway to regulate entry, with *ogt-1* acting upstream of the other genes (**Figure 1B; Figure 6**). On the other hand, while *sir-2.1* and *aak-2* also had defective entry, their phenotypes were distinct from the *ogt-1* pathway, particularly in relation to percentage of each fate adopted (**Supplemental Figure 2**) and fat storage (**Supplemental Figure 5**). Based on these distinct phenotypes, we postulate that *sir-2.1* and *aak-2* function in a distinct pathway to modulate ARD entry (**Figure 6**), and suggest that alternative metabolic mechanisms can act to regulate ARD entry.

**Figure 6.**
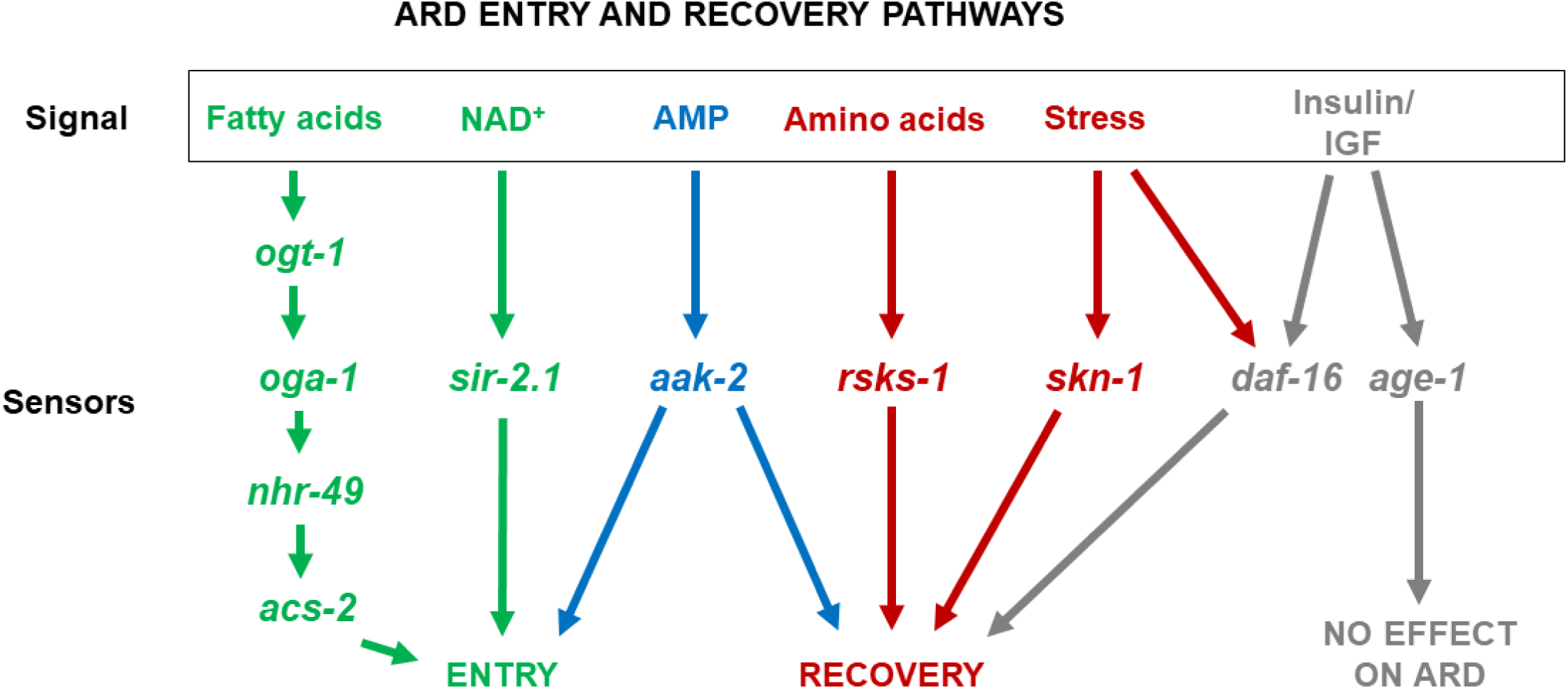
Model of genetic modifiers of ARD phases. Specific nutrient-sensing pathways contribute to ARD entry and exit. The initial sensing and dissemination of the signal to enter ARD (green pathways) involves genes encoding proteins that regulate fatty acid metabolism (*ogt-1, oga-1, nhr-49, acs-2*) and sirtuins (*sir-2.1*). Energy sensing, mediated by the product of *aak-2*, uniquely acts in both ARD initiation and recovery (blue pathway). The two members of the insulin/IGF signaling pathway tested in this study (grey pathways) showed differing results with *age-1* not affecting ARD, while the downstream *daf-16* affected recovery. ARD recovery additionally relies on the TOR pathway (*rsks-1*), which may also interact with *daf-16*. Stress signaling influences ARD recovery as well, with *skn-1* playing a minor role, and another possible pathway interacting with *daf-16*.

ARD maintenance, which was characterized by the shrinkage of the germline and retention of embryo(s) during ARD, was not significantly altered in any of the strains analyzed (**Figure 2**). Previous research found that apoptosis (CED-3) plays a role in maintenance (Angelo and Van Gilst 2009). Research with CED-3 and our results herein suggest that the maintenance of ARD may be independent of nutrient sensing but, rather, may rely more homeostatic processes coming into play during this extended period.

ARD recovery, as assessed by post-ARD lifespan, was disrupted in *aak-2, daf-16, rsks-1*, and, to a minor extent, *skn-1* mutants (**Figure 3**). While *daf-16* is most often associated with insulin-like signaling, it is also known to regulate and be regulated by the AMPK and TOR pathways (Antikainen *et al.* 2017). Based on the phenotypic similarity between *rsks-1* (TOR) and *daf-16*, we speculate that the TOR pathway is primarily responsible for recovery from ARD, particularly as loss of *age-1*, which acts upstream of *daf-16* in insulin-like signaling, did not have a significant impact on any of the ARD phases (**Figure 1B; Figure 2; Figure 3**). Further, while *aak-2* (AMPK) can also influence *daf-16* we find that *aak-2* had significant impact on both entry and exit, a distinct phenotype from *daf-16* (**Figure 1B; Figure 3**).

Overall, our findings support the previous assertion that ARD phases may be largely genetically separable (Angelo and Van Gilst 2009), with a general role for sensing changes in energy status both to enter and exit ARD. This complex constellation of regulators and distinct stages of ARD offers a promising model for future exploration of the contribution of these pathways in the regulation of longevity and may shed light on some of their dynamic roles under various pro-longevity regimens.

### O-GlcNAc cycling enzymes are required for initiation of ARD in concert with NHR-49

O-GlcNAc cycling is a highly dynamic nutrient-sensing post-translational modification (PTM) that allows for a rapid and reversible response to metabolic fluctuations. This PTM is uniquely situated at the node of multiple pro-longevity pathways and, thus, represents a promising target for therapeutic intervention (Love *et al.* 2010; Bond and Hanover 2014). However, the role of O-GlcNAc in longevity has yet to be fully explored, as it is essential in most organisms. Our results demonstrate that *ogt-1* and *oga-1* function in the sensing and dissemination of specific signals to enable ARD entry. Further, *ogt-1* and *oga-1* act in concert with the gene encoding the transcription factor NHR-49 to modulate ARD initiation (**Figure 1B**). Analysis of multiple alleles suggest that both *ogt-1* and *oga-1* are required for ARD entry but *oga-1* mutants are more variable in their effect on ARD entry. Biochemically, OGA-1 is known to act downstream of OGT-1 by dynamically removing the O-GlcNAc modification from proteins (Vocadlo 2012). Depending upon nutrient status, O-GlcNAc levels can vary widely with variable levels accumulating on proteins upon blocking O-GlcNAc cycling. It is also possible that OGA-1, while still functioning in the same pathway as OGT-1, may also have additional functions independent of O-GlcNAc removal. For example, it has been suggested that the *lin-4* microRNA controls the levels of both *sbp-1* and *oga-1* to regulate fat metabolism, longevity, ROS production and locomotion (Zhu *et al.* 2010).

OGA-1 has an evolutionarily conserved acetyltransferase-like domain whose precise function is not yet known (Toleman *et al.* 2004; Alonso *et al.* 2014). Both OGT-1 and OGA-1 are known to physically interact with several proteins and can form complexes that alter the localization and function of other proteins (Yang *et al.* 2002; Chen *et al.* 2013; Resto *et al.* 2016; Zhang *et al.* 2016; Eustice *et al.* 2017). While most of this information is derived from studies on mammalian O-GlcNAcase, the protein is highly conserved. Among the proteins that O-GlcNAcase binds are the fatty acid synthetases under conditions of stress (Groves *et al.* 2017) and OGA accumulates on lipid droplets in a manner suggesting a role in their assembly and mobilization (Keembiyehetty *et al.* 2011). Thus, OGA-1 may have other non-O-GlcNAc cycling functions, such as physically interacting with proteins in a way that suppresses, or otherwise interferes with, their activity. Indeed, our epistasis analysis of the role of *oga-1* in ARD entry suggests that OGA-1 may still functions biochemically in concert with OGT-1 to regulate NHR-49, but the regulation of fat metabolism may be more complex leading to a less penetrant defect in ARD entry.

Our epistasis analysis using multiple alleles of *ogt-1* and *oga-1*, suggests that *nhr-49, ogt-1*, and *oga-1* act in the same pathway (**Figure 1B**). Further, the ARD entry defects of *ogt-1* and *nhr-49* strains correlates with changes in fat stores (**Figure 5A**) but not carbohydrate stores (**Supplemental Figure 6**). Thus, our genetic and staining results suggest that modulation of fat stores is a key factor in ARD entry dynamics in the *ogt-1-*mediated ARD entry pathway.

NHR-49 regulates fatty acid metabolism by triggering fatty acid β-oxidation and/or desaturation (Van Gilst *et al.* 2005a). Several desaturases and unsaturated fatty acids are essential under different pro-longevity regimens, with multiple lines of evidence supporting an important role for lipid metabolism in pro-longevity pathways (Wang *et al.* 2008; Lapierre and Hansen 2012; Bustos and Partridge 2017). NHR-49 has a broad role in longevity, such that worms with impaired NHR-49 have decreased longevity under normal, fed conditions. This reduced lifespan under standard conditions is due to decreased fatty acid desaturation (Van Gilst *et al.* 2005a). On the other hand, both desaturation and β-oxidation of fatty acids mediated by NHR-49 is important for the increased longevity related to loss of the germline (Ratnappan *et al.* 2014). Our results demonstrate that ARD entry requires the β-oxidation pathway (*acs-2*) downstream of *nhr-49*. Overall, the regulation of fatty acid metabolism by NHR-49 has important implications for longevity, the mechanism of which may be context-dependent and dynamic.

Our analysis of the genes encoding the O-GlcNAc enzymes and *nhr-49* suggests a complex dynamic is at play. One interpretation of the findings is that O-GlcNAc cycling may act as a metabolic rheostat acting on NHR-49, thereby regulating the switch between fatty acid oxidation and desaturation, depending on environmental conditions. Previous work (Van Gilst *et al.* 2005a; Van Gilst *et al*. 2005b) has indicated that during starvation, mitochondrial β-oxidation (*acs-2* and other genes) increases fat consumption at the expense of *fat-7*-dependent fatty acid desaturation. We have also previously examined *fat-7* and *acs-*2 expression in the context of the *ogt-1* and *oga-1* mutants and their role in innate immunity and starvation where deregulation of both of these genes was observed. In this same paper, we also reported no change in the transcription of *nhr-49* for *ogt-1* and *oga-1* in L4 worms (Bond *et al.* 2014). These gene expression findings fit our genetic evidence as lipid stores are depleted in strains defective in ARD entry and *acs-2* seems to be critical for this utilization of stored fats under these conditions, whereas the *fat-7* mutants did not alter ARD initiation (**Figure 4B**).

### Potential Conservation of NHR-49/PPAR regulation of metabolism in mammalian torpor and hibernation

Recent findings suggest that the NHR-49 pathway in *C. elegans* shares many characteristics with PPARα in the mouse (Atherton *et al.* 2008; Taubert *et al.* 2006; Brandstädt *et al.* 2013). In mammals, PPARα is modified by O-GlcNAc which serves to modifiy its transcriptional activity (Ji *et al.* 2012). Intriguingly, PPARα also plays a key role in the control of torpor and FGF-21/NPY pathway in animals like hummingbirds, bats and hamsters (Ishida 2009). Torpor is a state of decreased physiological activity in an animal, usually characterized by a reduced body temperature and rate of metabolism. Torpor is often used to help these animals survive during periods of winter season, and it considered as the state in between sleep and hibernation. While there is a clear distinction between hibernation, torpor and diapause, the regulation of these processes may share some genetic determinants. Thus, the genetic findings reported here may potentially highlight a conserved role for O-GlcNAcylation in the NHR-49/PPAR pathway in torpor and hibernation. Indeed, recent findings in ground squirrels suggest that hibernation is associated with elevated O-GlcNAcylation of liver proteins (Jariwala *et al.* 2020).

### Summary and Conclusions

In this study, we have placed O-GlcNAc cycling in the context of other well established pro-longevity-associated nutrient sensors in modulating adult reproductive diapause. We have identified several novel regulators of ARD initiation including *sir-2.1, aak-2, acs-2, ogt-1* and *oga-1*, in addition to the previously described role for *nhr-49.* Further, we have shown that *skn-1, rsks-1*, and *daf-16* are important for ARD recovery. Defining key metabolites that are altered will provide additional insight into the central components regulated by the O-GlcNAc node and how these are altered under specific environmental conditions. Along with our novel genetic interactions with *nhr-49*, future exploration of the O-GlcNAc cycling enzymes interactions with nuclear hormone receptors may provide new insight into the role of OGT and OGA as metabolic rheostats. Beyond impaired ARD entry, we found that loss of the O-GlcNAc cycling enzymes converges on the fatty acid metabolic pathways involving mitochondrial fatty acid β-oxidation. The findings described herein indicate that ARD entry and recovery are largely genetically separable and that O-GlcNAc cycling acts with other nutrient sensors in ARD initiation. These results have important implications for understanding the dynamic activity of nutrients sensors during DR and for defining the role of these sensors in regulation of mammalian lipid mobilization and metabolism.

## ACKNOWLEDGMENTS

The authors wish to thank the members of the Hanover and Krause labs and the Baltimore-Washington Worm Club for helpful discussion and input on this manuscript. We also acknowledge the *C. elegans* Genetics Consortium for the many *C. elegans* strains provided by this resource. We would like to thank Marc Van Gilst for supplying the original *nhr-49* allele used in the study. The study was funded by intramural NIDDK research funding which is gratefully acknowledged.

## SUPPLEMENTAL FIGURE AND TABLE LEGENDS

**Supplemental Figure 1. Summary of ARD protocol**

As described in Materials and Methods, worms were bleached to obtain synchronized L1 larvae. The L1s were then plated at a density of 30,000 per plate, harvested at mid-L4 stage, and grown on plates without food for 48-72 hours. At this stage worms were counted to determine the percentage of animals in arrested L4, bagged adults, or ARD (arrows indicate arrested embryos) as shown in the representative photos.

**Supplemental Figure 2. Fates adopted by each strain during initiation of ARD**.

The L4 and bagging fates tended to have a higher degree of variability than ARD entry, as evidenced by a higher SEM. Select strains had a marked increase in the percentage of worms in L4 including *aak-2(ok524), age-1(hx546), daf-16(mu86), sir-2.1(ok434)*, and *skn-1(zj15).* Other mutant strains, *nhr-49(nr2041), oga-1(ok1207)*, and *ogt-1(jah01)*, were more similar to wildtype in terms of having a more even distribution between L4 and bagging even though this set was defective for ARD entry. It is worth noting that *oga-1(ok1207)* and the full deletion allele *oga-1(av82)* displayed a significant defect in ARD entry. However, another allele, *oga-1(tm3642)* exhibited a significantly stronger defect in ARD entry. Considering the heavy mutational load used to create high throughput alleles, the stronger phenotype of *oga-1(tm3642)* may be due to a linked background mutation, as has been recently described for an allele from the same source (Bauer *et al.* 2019). As such, we chose to exclude this allele from further genetic analysis.

**Supplementary Table 1. ARD fates.**

Table representation of Supplemental Figure 2. Average percentage of each strain adopting each fate with SEM shown to illustrate variability.

**Supplemental Table 2. Brood size variability.**

The table shows the average brood size under normal husbandry (control) or re-fed animals following 5 or 10 days in ARD. A high degree of variability was observed between and within experimental replicates for the average brood size (SEM). These brood sizes did not correlate to fate adoption of each strain nor with influence on any of stages of ARD. Red indicates a decrease compared to wildtype whereas green indicates an increase vs wildtype.

**Supplemental Figure 3. DAPI staining of spermatheca during ARD reveal changes in number of sperm nuclei during ARD.**

(A) DAPI staining of worms after 15 days in ARD reveals fewer sperm in the spermatheca of *ogt-1(1474)*, and *nhr-49(nr2041)* compared to wild-type N2 worms. In the *oga-1(ok1207)* worms, more sperm were observed. White arrows indicate examples of sperm nuclei. (B) Whereas most of the strains analyzed saw a decrease in the number of sperm nuclei present at day 30 of ARD, *oga-1(ok1207)* had an overall increase compared to wildtype. However, these dynamics did not correlate with changes in brood size among selfed individuals. Green arrows indicate increased brood numbers vs wildtype and red arrows indicate decreased brood vs wildtype.

**Supplemental Figure 4. 24 hours of ARD results in noticeable decrease in ORO staining.**

(A) Decrease in ORO signal were observable at 24 hr after worms were placed on ARD places, as shown with representative strains.

(B) Strains were stained with DAPI to demonstrate efficient small-molecule penetrance of cuticles across strains, indicating that changes in ORO staining are not related to differences in cuticle penetrance between strains.

**Supplemental Figure 5. Additional strains stained for ORO after 30 days of ARD.**

In addition to the *ogt-1-*dependent pathway (shown in figure 5) we also looked at strains with diverse phenotypes to see if changes were unique to the *ogt-1* pathway. We observed that outside of the *ogt-1* pathway, strains that did not influence either entry or exit (*age-1(hx546)*), strains that also only influenced entry (*sir-2.1(ok424)*), and strains that influenced only recovery (*skn-1(zj15)*) had a marked increase in ORO staining/TAG stores compared to both wildtype and the *ogt-1* pathway. We also observed this same pattern with genes downstream of *nhr-49*, such that *acs-2* (with a defective ARD entry) had reduced ORO staining but *fat-7* (no defect in ARD entry) had a stronger ORO signal. Arrows indicate arrested embryos, which show high ORO staining.

**Supplemental Figure 6. Changes in glycogen and trehalose levels before and after ARD.**

(A) Carminic acid staining (indicative of glycogen and trehalose levels) varied greatly between strains. As we have previously reported (Hanover *et al.* 2005), *ogt-1(1474)* showed slightly higher levels of carminic acid staining than other strains in standard husbandry conditions, though in this study this change did not reach significance. After 30 days of ARD (lower panels) we noted that *nhr-49(nr2041)* had a dramatic increase in staining, while wild type and *ogt-1(ok1474)* did not. These results did not correlate with the observed defect in ARD entry for these strains. ImageJ based quantification of carminic acid fluorescence by pixel intensity. *P*-value **** = <0.0001, as determined by two-way ANOVA.

